# Difference in Differences analysis evaluates the effects of the Badger Control Policy on Bovine Tuberculosis in England

**DOI:** 10.1101/2023.09.04.556191

**Authors:** Colin P. D. Birch, Mayur Bakrania, Alison Prosser, Dan Brown, Susan M. Withenshaw, Sara H. Downs

**Affiliations:** Animal and Plant Health Agency, Woodham Lane, Addlestone, Surrey UK, KT15 3NB

## Abstract

Persistent tuberculosis (TB) in cattle populations in England has been associated with an exchange of infection with badgers *(Meles meles*). A badger control policy (BCP) commenced in 2013. Its aim was to decrease TB in cattle by reducing the badger population available to provide a wildlife reservoir for bovine TB. Monitoring data from 52 BCP intervention areas 200-1600 km^2^ in size, starting over several years, were used to estimate the change in TB incidence rate in cattle herds, which was associated with time since the start of the BCP in each area. A Difference in Differences analysis addressed the non-random selection and starting sequence of the areas. Herd incidence rate of TB reduced by 56% (95% Confidence Interval 43-67%) up to the fourth year of BCP interventions, with the largest reductions in the second and third years. There was insufficient evidence to judge whether incidence rate reduced further beyond four years. These estimates are the most precise for the timing of decline in cattle TB associated with interventions primarily targeting badgers. They are within the range of previous estimates from England and Ireland. This analysis indicates the importance of reducing transmission from badgers to reduce the incidence of TB in cattle, noting that vaccination of badgers, fertility control and on farm biosecurity may also achieve this effect.

## Introduction

Bovine tuberculosis (TB) is an important problem for the cattle industry in the UK, associated with substantial economic costs, implications for trade and risks to animal and human health. It is an infectious, zoonotic bacterial disease, caused by *Mycobacterium bovis*, which infects a wide range of animals ^[1-3]^. Eradicating TB from the British cattle population requires a combination of approaches ^[4,5]^. The disease is difficult to eradicate from domesticated animal populations when there is a local reservoir of infection in wildlife, without removing the infection in wildlife ^[6-8]^.

Evidence of exchange of *M. bovis* infection between cattle and badgers suggests that coordinated TB control in both species may be necessary to control infection in cattle ^[9,10]^. Badger culling as an intervention to reduce TB incidence in cattle has been implemented at different times and at defined locations within England since the 1970s ^[6,11,12]^. However, the impact of badger culling is contentious ^[13-15]^. The most thorough study in England of the effect of badger culling on TB in cattle was the Randomised Badger Culling Trial (RBCT) conducted between 1998 and 2005. Incidence of confirmed TB in cattle herds was overall c. 29% (95%CI 21 - 36%) lower in areas with widespread systematic culling than in non-intervention areas ^[16]^. However, interpretation of the RBCT as evidence for or against badger culling has been disputed ^[17]^. A badger control policy (BCP) with licensed culling of the European badger *(Meles meles)* commenced in England in 2013 ^[18]^. Reducing badger population density may lower TB prevalence in badgers as well as encounters between badgers and cattle ^[19]^. This policy has been supported by associated measures including the provision of biosecurity advice, as well as additional testing of cattle herds under restriction to manage TB incidents. Descriptive data including the incidence and prevalence of bovine TB in areas subject to BCP controls have been published annually since 2014 ^[20]^.

By the end of 2020, in regions of England designated as having relatively high incidence of bovine TB, 52 areas of various sizes (200 – 1600 km^2^) were issued with licences to cull badgers. BCP controls started in different years in different areas, from 2013 to 2020 ^[20]^. Previous analysis of the effects of the BCP by comparing cattle TB incidence in the first three licensed BCP areas with incidence in matched control areas, whilst also controlling for confounding factors, estimated a reduction in herd incidence rates of between 37-66% ^[21,22]^. However, the increase in the assignment of new land to the BCP reduced the availability of comparable unculled land, preventing adequate matching to control areas. A different approach was required. Simple retrospective comparison of BCP areas with unmatched non-culled areas would be misleading because of confounding bias ^[23]^. Farmers applied for licences voluntarily, and areas were required to meet several conditions for a badger culling licence to be granted. As such, the BCP areas were not chosen at random, so their background herd density and levels of bovine TB differed from non-culled areas. We therefore undertook a new approach, which became feasible with the large number of areas assigned to the BCP. We compared incidence rates within and between BCP areas using a Difference in Differences analysis ^[24]^, with the areas defined by BCP badger culling licences being the units in the analysis.

## METHODS

### Data

Each BCP area was selected by the extent of support from local landowners to form a company to carry out the culling of badgers and their capability to adhere to licence criteria. Its boundary was then defined in a licence agreement with Natural England, the national regulator. For this study, boundary information was used for the 52 BCP areas located in relatively high bovine TB incidence regions in England (the “High Risk Area” and “Edge

Area”), where culling began between 2013 and 2020 ^[20]^. (Badger culling licences were also obtained for two relatively small areas in the Low Risk Area of England, which were not comparable with the areas included in this analysis.) Data were obtained on the cattle herds and bovine TB incidents within the 52 areas from 1^st^ September 2009 to 31^st^ December 2021, from routine surveillance data held on APHA’s bovine TB management database. The BCP was rolled out in a phased manner, as BCP interventions were started over a series of years in 1 to 11 areas each year. As a result, in any year there were various numbers of areas that had been exposed to BCP controls for different lengths of time (Table 1). For example, by December 2020, 21 areas had been subject to 4 or more years of the BCP control.

**Table 1:**
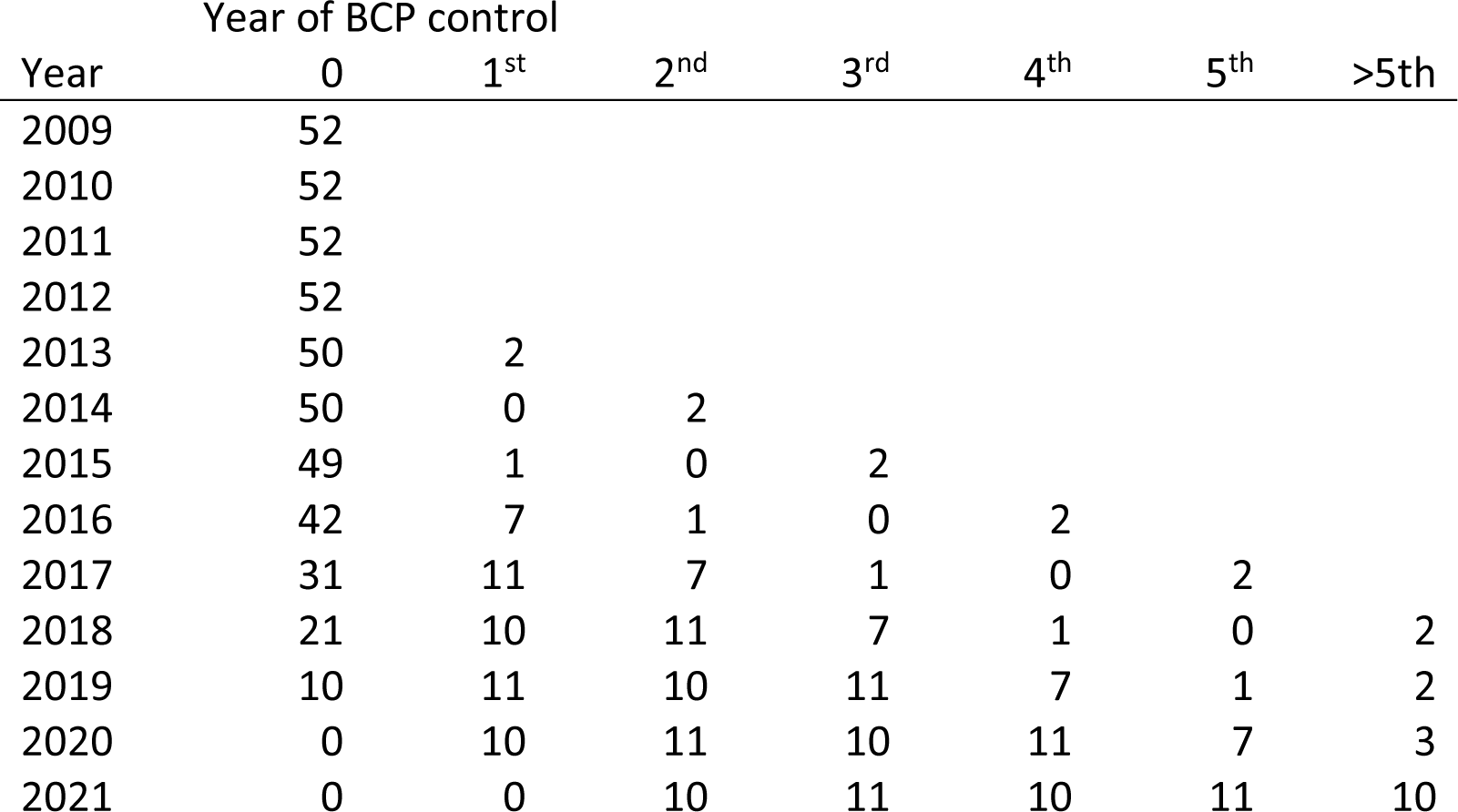
Numbers of areas subject to the Badger Control Policy (BCP) in December of each year, 2009-2021. The data ended at the end of December 2021, so the last intervention year started in 2020.

| Year | Year of BCP control |  |  |  |  |  |  |
| --- | --- | --- | --- | --- | --- | --- | --- |
|  | 0 | 1<sup>st</sup> | 2<sup>nd</sup> | 3<sup>rd</sup> | 4<sup>th</sup> | 5<sup>th</sup> | >5<sup>th</sup> |
| 2009 | 52 |  |  |  |  |  |  |
| 2010 | 52 |  |  |  |  |  |  |
| 2011 | 52 |  |  |  |  |  |  |
| 2012 | 52 |  |  |  |  |  |  |
| 2013 | 50 | 2 |  |  |  |  |  |
| 2014 | 50 | 0 | 2 |  |  |  |  |
| 2015 | 49 | 1 | 0 | 2 |  |  |  |
| 2016 | 42 | 7 | 1 | 0 | 2 |  |  |
| 2017 | 31 | 11 | 7 | 1 | 0 | 2 |  |
| 2018 | 21 | 10 | 11 | 7 | 1 | 0 | 2 |
| 2019 | 10 | 11 | 10 | 11 | 7 | 1 | 2 |
| 2020 | 0 | 10 | 11 | 10 | 11 | 7 | 3 |
| 2021 | 0 | 0 | 10 | 11 | 10 | 11 | 10 |

The number of herd incidents in each area each year was standardised by the herd time at risk (herd years) to generate incidence rates ^[20]^. For the primary analysis, herd incidents were restricted to those confirmed by visible tuberculous lesions in one or more TB test-positive animals (reactors) removed from the herd, or the identification of *Mycobacterium bovis* by bacteriological culture (herds with “Official Tuberculosis Free status Withdrawn” (OTFW)). Annual units were intervention years starting on 1^st^ September and ending 31^st^ August the following year, so their start roughly matched the timing of annual badger culls which were scheduled to run for 6 weeks from near the beginning of September. The intervention year defined the number of badger culls already completed in each area. The herd population included all active herds present within the defined boundaries of each area on 1^st^ September each year (“Herds in Existence” (HIE)). A minority of BCP areas had minor boundary extensions over time. For each area, HIE was defined using the most recently defined area boundaries up to 2021 ^[20]^.

Data were also available for all bovine TB incidents, including “OTF suspended” (OTFS) incidents, where reactors to the Single Intradermal Comparative Cervical Tuberculin (SICCT) test had been detected in a herd but infection had not been confirmed by the presence of visible lesions or culture of *M. bovis*. Overall, OTFS incidents were 28.5% of all incidents (i.e. OTFS + OTFW) but almost all false positive incidents were OTFS. Opportunities to confirm an incident as OTFW increased with the number of TB test reactors in the herd, so true positive OTFS incidents were likely to include fewer reactors on average than OTFW incidents. Some factors associated with the probability that a case was confirmed were also associated with the probability of detection, e.g. recently infected cattle would be less likely to react to the SICCT test and less likely to have lesions ^[25]^. Thus, the number of OTFS incidents was more sensitive to changes in the sensitivity of surveillance than was the number of OTFW incidents. Changes to surveillance potentially obscured changes to the true burden of disease. Therefore, as in previous analyses of the effect of badger culling ^[16,21]^, OTFW incidence rate was felt to be a more reliable indicator of changes in the true level of TB infection in cattle than the total (OTFW + OTFS) incidence rate and so was the outcome variable of the primary analysis.

Calendar year is a more familiar annual unit than intervention year. However, each calendar year when badger controls started only included a period of 4 months following the start of interventions. Only 1/3 annually tested herds would expect to be tested during this time and a detectable effect on cattle TB incidence rate so soon after badger culling was unlikely. Therefore, the intervention year provided a clearer contrast between the last year before controls started and the first year after, and so was used as the annual unit in the primary analysis.

Over time, some herds that were present in a BCP area when controls were first introduced (i.e. the original cohort of herds) may have ceased operation, while new herds may have come into existence. New herds might be more likely to bring in cattle that had not been exposed to the local controls, and so potentially reduce power to detect an effect of those controls. However, since herds that cease operation are not a random selection from the herd population, HIE was judged to be a preferable study population for the primary analysis.

For comparison, in addition to the primary analysis, alternative analyses were completed for all eight combinations of OTFW vs all incidents, intervention year vs calendar year and incidents in HIE vs only cohort herds. For each area, the cohort was defined each year as the subset of the original cohort that remained active at the start of the year.

### Analysis

Bovine TB incidence rates in BCP areas were summarised by year to show the overall changes from the effects of BCP interventions interacting with underlying trends. These were compared with summaries of incidence rates in areas while not under BCP controls, to show trends without the effects of interventions. This summary of the observations with minimal analysis provided a check on the outputs from more formal analysis.

The impact of BCP controls on bovine TB was inferred by a Difference in Differences method ^[24]^. A statistical model was fitted to the incidence rates observed in each of the 52 BCP areas each year during 2009-2021. The model included fixed effects for differences in average incidence between BCP areas and years. It also fitted a local linear trend of incidence rate within each area up to the start of badger controls. This early linear trend had two purposes. It could improve the estimate of the level of bovine TB before the start of badger culling by allowing an influence from earlier years while giving higher weight to the years closer to the start of badger culling. It also matched differences between areas in the trends of bovine TB before the start of TB culling, which included a trend for variation in herd incidence rate between areas to decrease over time.

When analysing by intervention year, we treated each year of BCP control up to the 4^th^ intervention year as a distinct stage of control, with years beyond this merged as a single last stage, so the BCP factor had 6 levels: before BCP, the four intervention years 1-4 and the fifth and later intervention years merged (Table 1). When analysing by calendar year, the BCP factor had 7 levels: before BCP, the first calendar year of BCP, which included less than 4 months after the first badger cull, the four calendar years 2-5 of BCP and later years. Observations ended at the end of December 2021, so the numbers of observations of calendar years 2-5 and 6+ of BCP matched intervention years 1-4 and 5+ (Table 1).

Counts of disease incidents are often analysed by a Generalised Linear Model (GLM) using Poisson regression ^[21]^, but the data reported here have been regularly reported as incidence rates ^[20]^. Incidence rates allow transparent comparison of burden of disease between areas with widely different sizes; they can also correct for differences in surveillance intervals and restrictions between areas and over time. A linear analysis after square-root transformation is an alternative approach suitable for an analysis of incidence rate, which also relaxes the assumptions about the relationship between mean and variance implied by Poisson regression. Therefore, primary analysis was by linear regression of square-root transformed incidence rates, which also allowed easier interpretation and exploration of the common trend assumptions of the Difference in Differences design ^[24]^. The number of incidents was increased by 0.5 before calculating the transformed incidence rate to mitigate the influence of a few observations of zero incidents ^[26]^. Examination of square-root transformed observations from areas before BCP controls started locally confirmed transformation improved homogeneity of variance and linearity of data.

Thus, the linear statistical regression model with square-root transformed incidence rate as the dependent variable was:

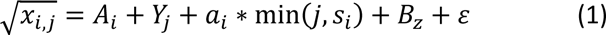

Where *x_i,j_* = incidence rate in area *i* during year *j*, *A_i_* = a coefficient shared by incidence rates in area *i*, *Y_j_* = a coefficient shared by incidence rates during year *j*, *a_i_* = an annual change of incidence rates in area *i* up to the year before BCP controls began in the area (*s_i_* = the year before BCP controls began), *B_z_* = a coefficient shared by areas in stage *z* of badger control, given that area *i* during year *j* is at badger control stage *z*, while *ε* = an error term. In the model (1) there was some redundancy among the parameters *A_i_*, *Y_j_*, *B_z_* and *a_i_*. The average value of *a_i_* effectively adjusts the values of the other parameters. This redundancy was addressed by constraining the sum of *a_i_* to zero. This constraint ensured that *a_i_* was fitted to differences in trends among areas before the start of BCP interventions, while *B_z_* was fitted to differences in trends among areas after the start of BCP interventions. The model’s estimate of an area’s incidence rate in the year before BCP started was the baseline setting the incidence rate expected with no BCP effect in later years. The analysis was executed using the constrained regression “cnsreg” command in Stata^(R)^ 15.0 (StataCorp). Linear predictions to illustrate model outputs were generated using the postestimation “margins” command in Stata^(R)^ 15.0 (StataCorp).

All alternative data sets were also analysed using an equivalent Poisson regression, i.e. an analysis of counts (number of incidents per area per year) with a log link function, with herd time at risk included as an exposure variable. This regression was also run with the sum of *a_i_* constrained to zero, using the “poisson” command in Stata^(R)^ 15.0.

## RESULTS

### Trends in observed incidence rate

Trends in incidence rate were obscured by the relatively large variation in incidence rate between areas. Median OTFW incidence rate across all 52 areas rose to a maximum during 2013-2015 followed by a decrease to the end of the study (Fig. 1a). Note that 2016-2017 was the first year more than three areas were under BCP controls. By the 2018-2019 intervention year, the OTFW median incidence rate dropped below its value in 2009-2010 and continued to decline. The range of incidence rates reduced over time, which was a major reason for including trends differing between areas before the start of badger culling. Badger culling did not start until 2013, so the early increase in incidence rate was also visible when only considering data from areas while they were not exposed to BCP controls (Fig. 1b, c). However, in contrast to the overall trend including areas under badger culling, median OTFW incidence rate in unculled areas did not substantially decline after 2015. For example, the cohort of areas that did not start controls until 2019-2020 had lower median incidence rate than all areas up to 2014-2015, but higher median incidence rate in 2018-2019 (Fig. 1c compared with Fig. 1a). Before 2013, the 2019-2020 cohort had lower median incidence rate than the cohort that started badger culling during 2013-2018, suggesting non-random selection of areas for controls (Fig. 1b, c). These observations provide informal evidence suggesting the trend across all areas was related to the introduction of BCP controls rather than other causes.

**Figure 1:**
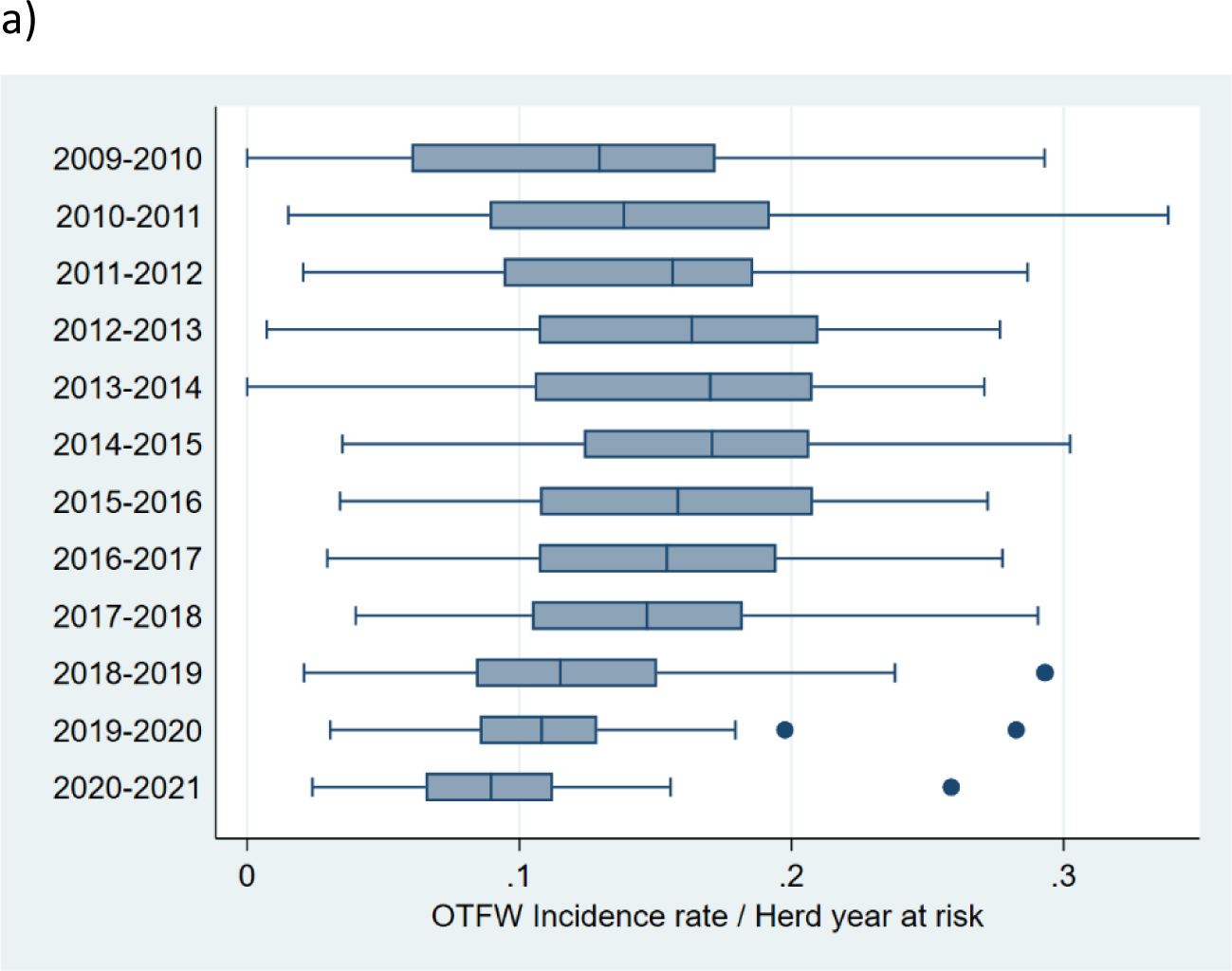
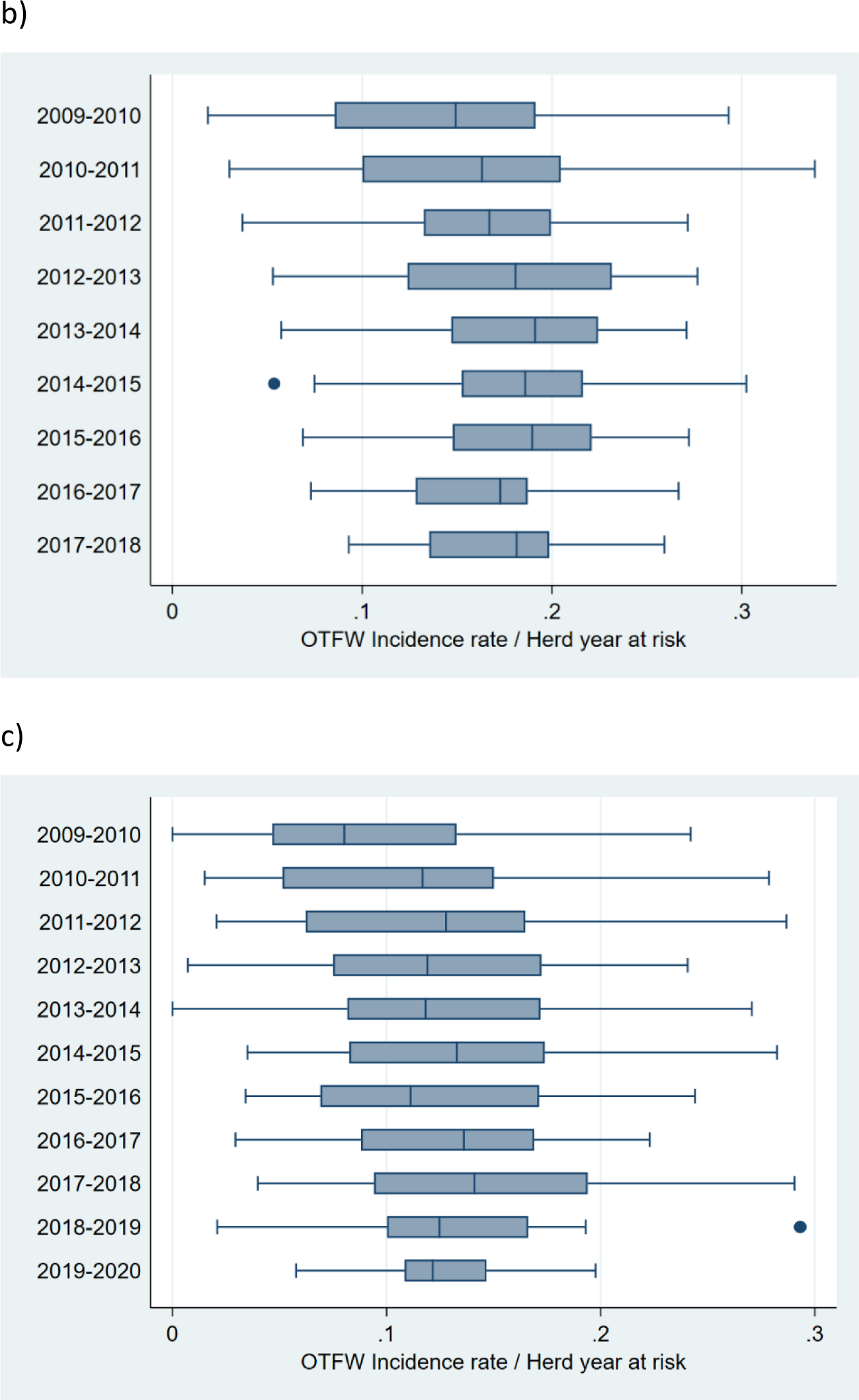
OTFW incidence rates (OTFW incidents per herd years at risk). a) All 52 BCP areas included for each intervention year from 2009 to 2021. b) 31 areas that started badger culling during 2013-2018; each area is only included before badger culling started, so the cohort gradually reduced from 2013 (see Table 1). c) 21 areas that started badger culling in 2019 (11) or 2020 (10). Only the last 10 areas are included for 2019-2020. In the box and whisker plots, the central vertical line is the median. The box ranges from the lower quartile to the upper quartile. The whiskers show the full range of the data, unless outliers are present, which are shown individually as dots.

The trend in OTFS incidence rate did not follow OTFW incidence rate, fluctuating between years in a more irregular way (Fig. 2). The median OTFS incidence rate was lowest in 2014-2015 but was higher in all years from 2015-2016 onwards than in all years before 2015.

**Figure 2:**
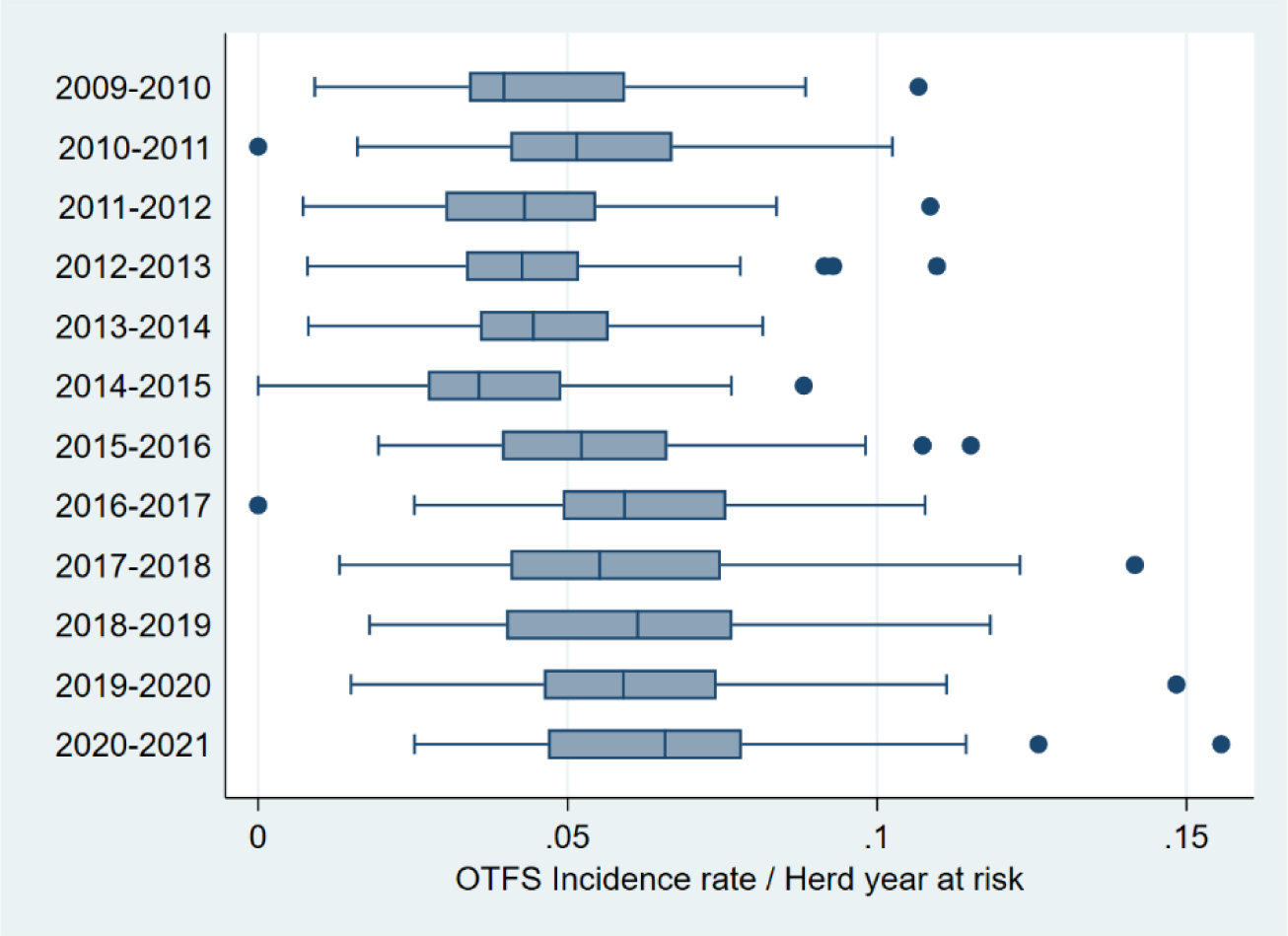
OTFS incidence rate (OTFS incidents per herd years at risk) for all 52 BCP areas for each intervention year from 2009 to 2021. See Fig. 1 for explanation of presentation.

### Area effects from the statistical model

Incidence rate differed substantially between areas and the differences were relatively consistent over time. Thus, although they were not of interest in themselves, the area effects were the largest component of the linear statistical model (Table A1 in Appendix). The incidence rates also tended to be lower in areas that started badger culling in later years (Fig. 3). The trend was much clearer from the outputs of the statistical model than from the trends in the raw data (Fig. 1), because the average estimated incidence rates across areas from the statistical model had narrow confidence intervals, which reflected the consistency of the incidence rate within each area (Fig. 3). The Difference in Differences analysis took account of consistent differences between areas to improve the precision of its estimate of BCP effects and reduce bias.

**Figure 3:**
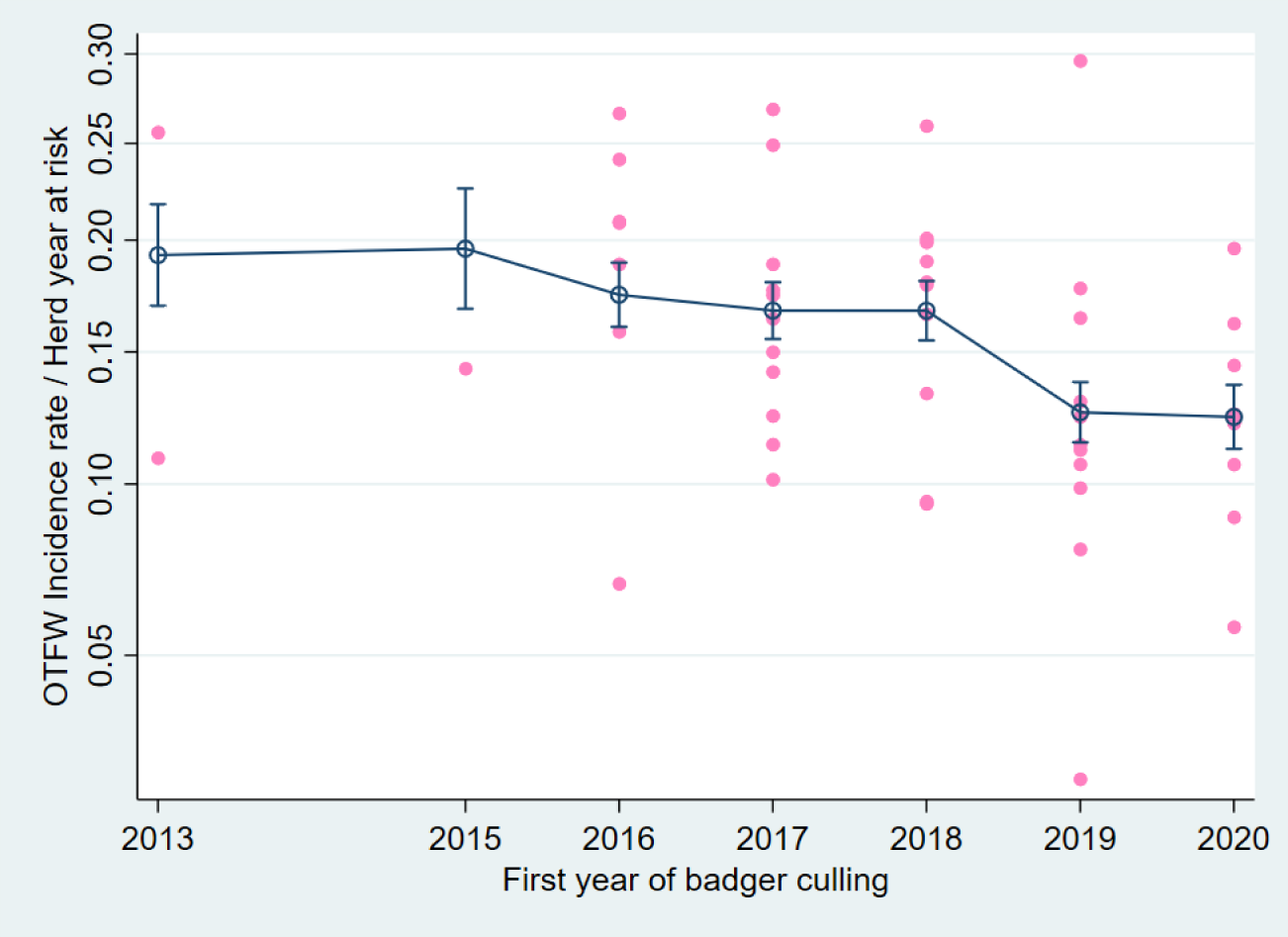
The OTFW incidence rate the year before starting badger culling related to the year culling started. Linear prediction from the statistical model (linked blue circles with 95% confidence intervals) over observed values (pink dots, one for each area). The scaling of the vertical axis is square-root transformed incidence rate, although it is labelled with incidence rate.

### The effects of BCP over time

Overall, the OTFW incidence rate decreased with the duration of badger culling (Fig. 4, Table 2). The incidence rate in the first intervention year of BCP decreased, although the reduction was relatively small (Fig. 4a). In the second and third intervention years of BCP the incidence rate decreased by larger amounts. The width of confidence intervals increased with time after the start of controls, so it was not clear whether and when the maximum effect of BCP interventions was achieved. However, the steepest declines in incidence rate were before the end of the third intervention year, and there appeared to be more decline beyond the third year. Analysis by calendar year found an equivalent trend (Fig. 4b). During the first calendar year, which only included the first 4 months after the start of culling, bovine TB herd incidence rate was similar to before controls. The second to fifth calendar years of BCP roughly matched the first to fourth intervention years. Across the 52 areas and 12 years, the expected incidence rate without BCP was approximately 0.145 incident per herd year at risk. In the fourth intervention year, the equivalent incidence rate was reduced to c. 0.065, a reduction of approximately 56% (Table 2).

**Figure 4:**
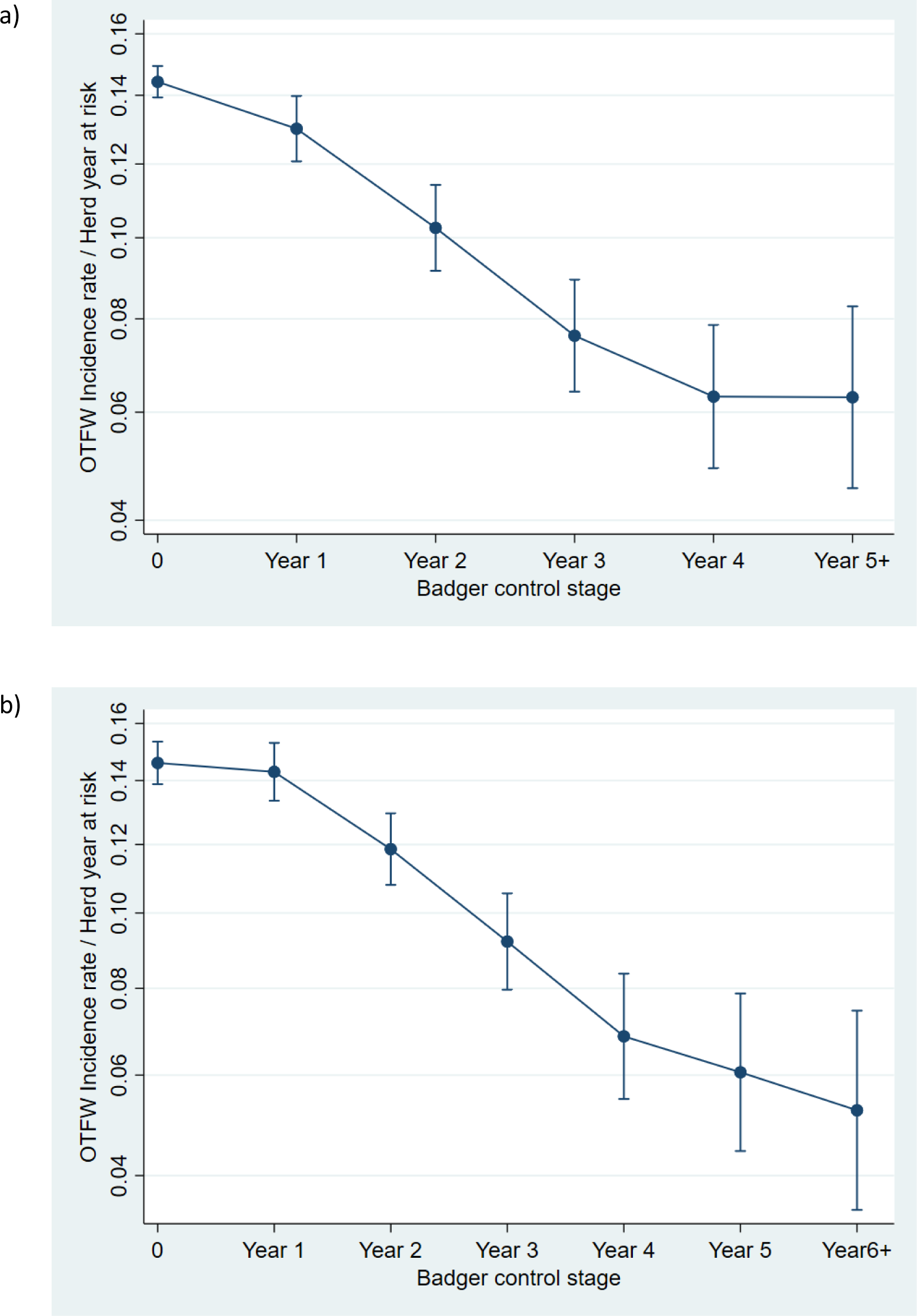
Linear predictions from the statistical model of confirmed (OTFW) incidence rate related to local duration of Badger control policy shown for a) intervention years (September to August) and b) calendar years. Error bars show 95% confidence intervals. The scaling of the vertical axis is square-root transformed incidence rate, although it is labelled with incidence rate.

**Table 2:** Percentage reduction of incidence rate associated with local duration of Badger control policy in intervention years.

| Intervention year | Estimated Reduction (%) | 95% Confidence interval (%) |
| --- | --- | --- |
| 1 | 9.8 | 1.2 – 18.0 |
| 2 | 28.7 | 18.9 – 37.9 |
| 3 | 46.8 | 35.9 – 56.6 |
| 4 | 55.8 | 43.4 – 66.7 |
| 5+ | 56.0 | 40.7 – 69.0 |

### Trend over time

Model outputs indicated that the potential trend in the absence of BCP interventions was relatively little decline of the incidence rate from its peak in 2014-2015, with the average OTFW incidence rate remaining within the range 14-16% from September 2012 to August 2021 (Fig. 5). This trend contrasted with the observed trend with BCP as applied (Fig. 1a), and was consistent with observations limited to areas that had not started badger culling (Fig. 1b, c). The overall decline of incidence rate after 2015 seems to have been mainly due to the effect of BCP.

**Figure 5:**
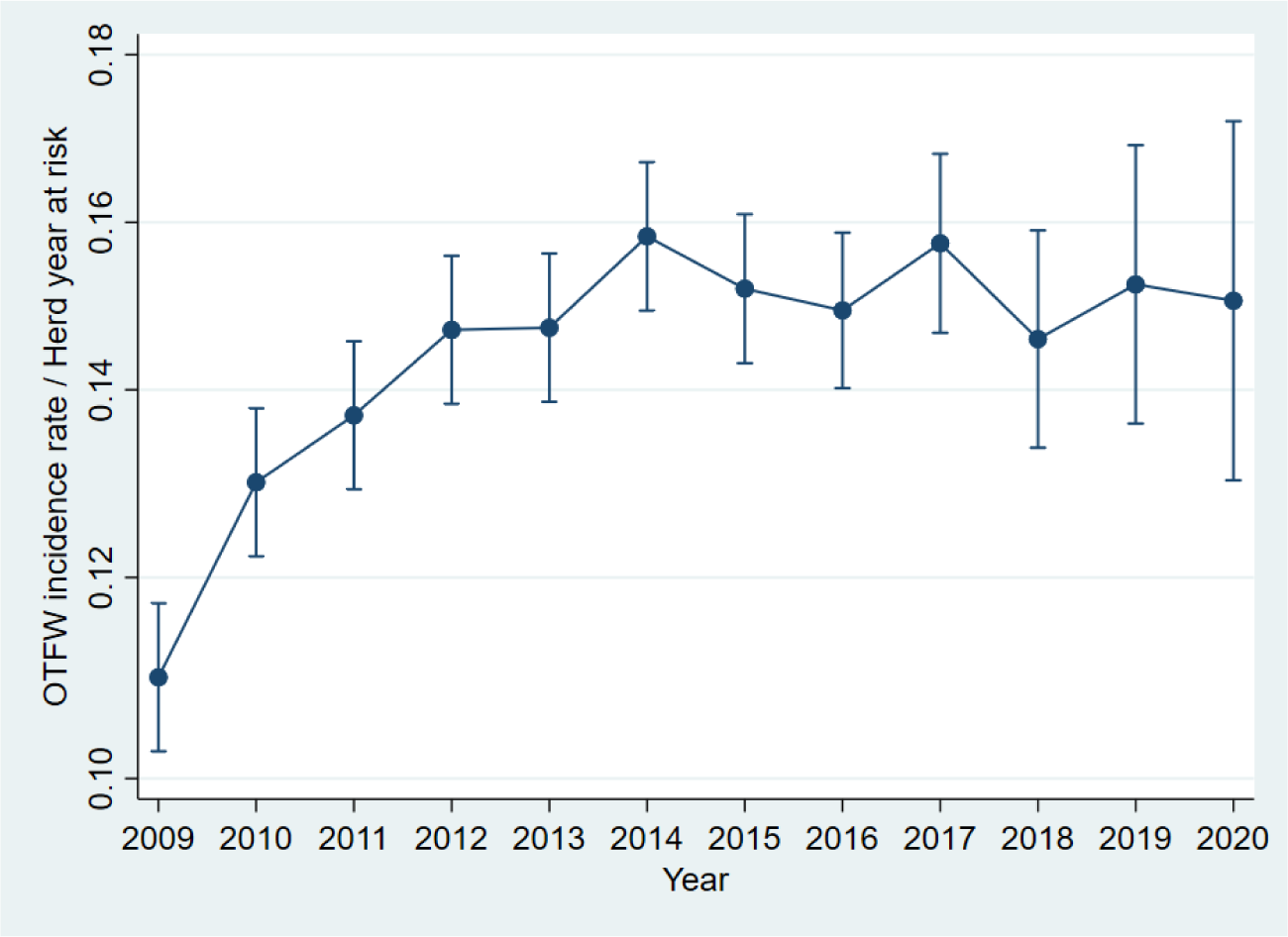
Average OTFW incidence rate for each intervention year observed or estimated without BCP interventions across the 52 BCP areas. Labels show when intervention years started. Vertical bars indicate 95% confidence intervals.

### Visual fit of model

The statistical model was compared with observed incidence rates in each area to allow visual review (Fig. 6). The multiplot illustrates the importance of the area effect: a horizontal line at a different level for each area would represent much of the observed variation.

**Figure 6:**
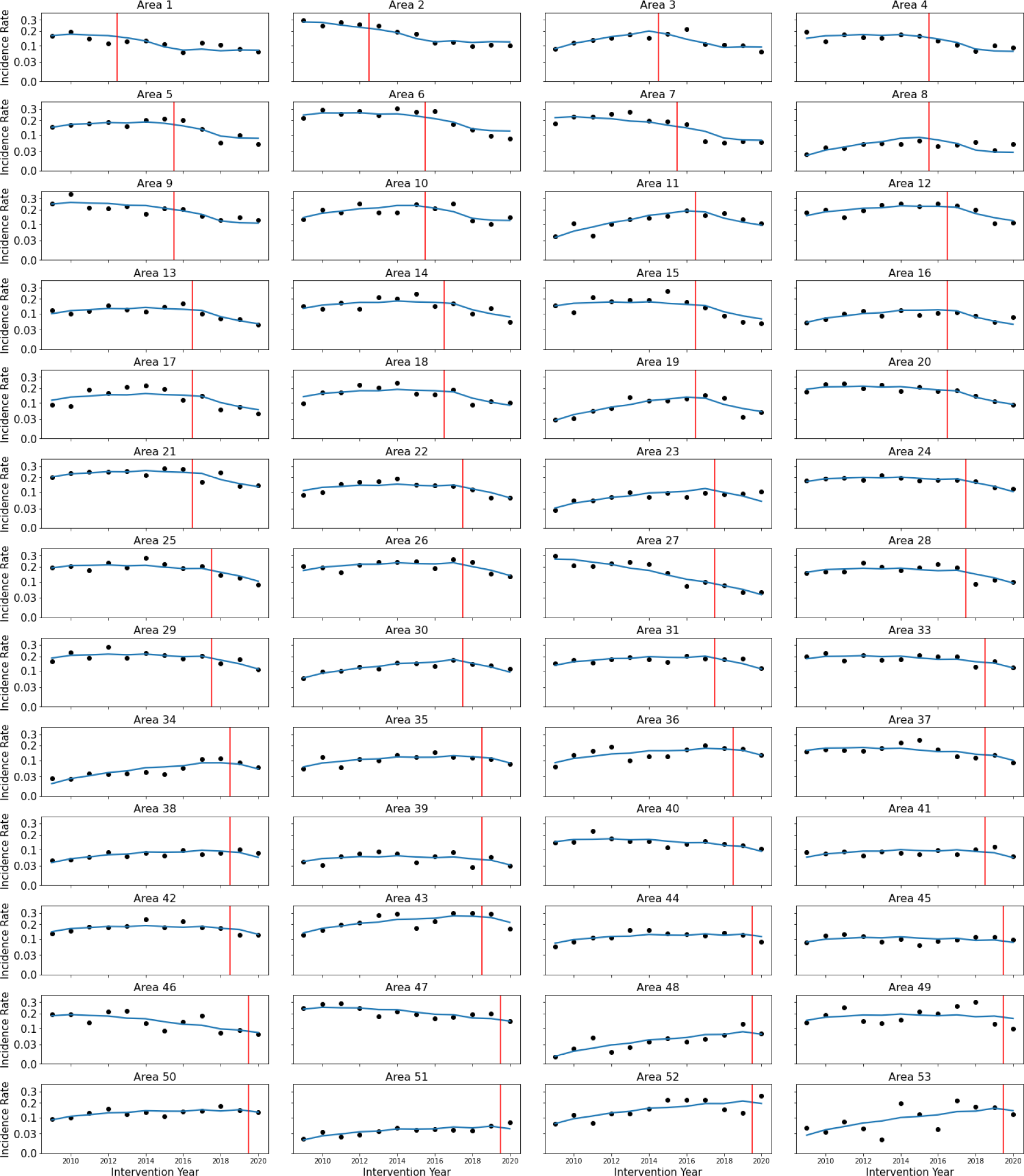
Comparison of the statistical model (blue line) to observed incidence rate i.e. OTFW per Herd year at risk (black dots) for each area. The red line represents the start of the BCP in that area. Area numbers follow the regular monitoring reports ^[20]^.

Nevertheless, the trend before the start of BCP clearly varied among areas. The analysis demonstrated significant variation (P < 0.0001) in the regression slopes up to the year before BCP started (Table A1 in Appendix). The fitted curves also emphasize the general trend for local incidence rate to decline in individual areas after BCP controls started. The model was a closer fit for some areas than others, partly because the incidence rate was observed with more uncertainty in smaller areas with fewer incidents. Across the areas, HIE ranged from about 40 to over 1000 herds. However, the model seemed broadly consistent with the observed differences between areas and local temporal trends.

### Trends in individual areas

Trends in incidence rate before BCP started differed between areas (Fig. 6). There was no evidence of a relationship between the year BCP started in an area and the trend of TB incidence rate before BCP started (Fig. 7). Most slopes for change of square-root transformed OTFW incidence rate had absolute value less than 0.02 yr^-1^, which would be equivalent to an annual change of about 10% from an incidence rate around 0.15, i.e. to

**Figure 7:**
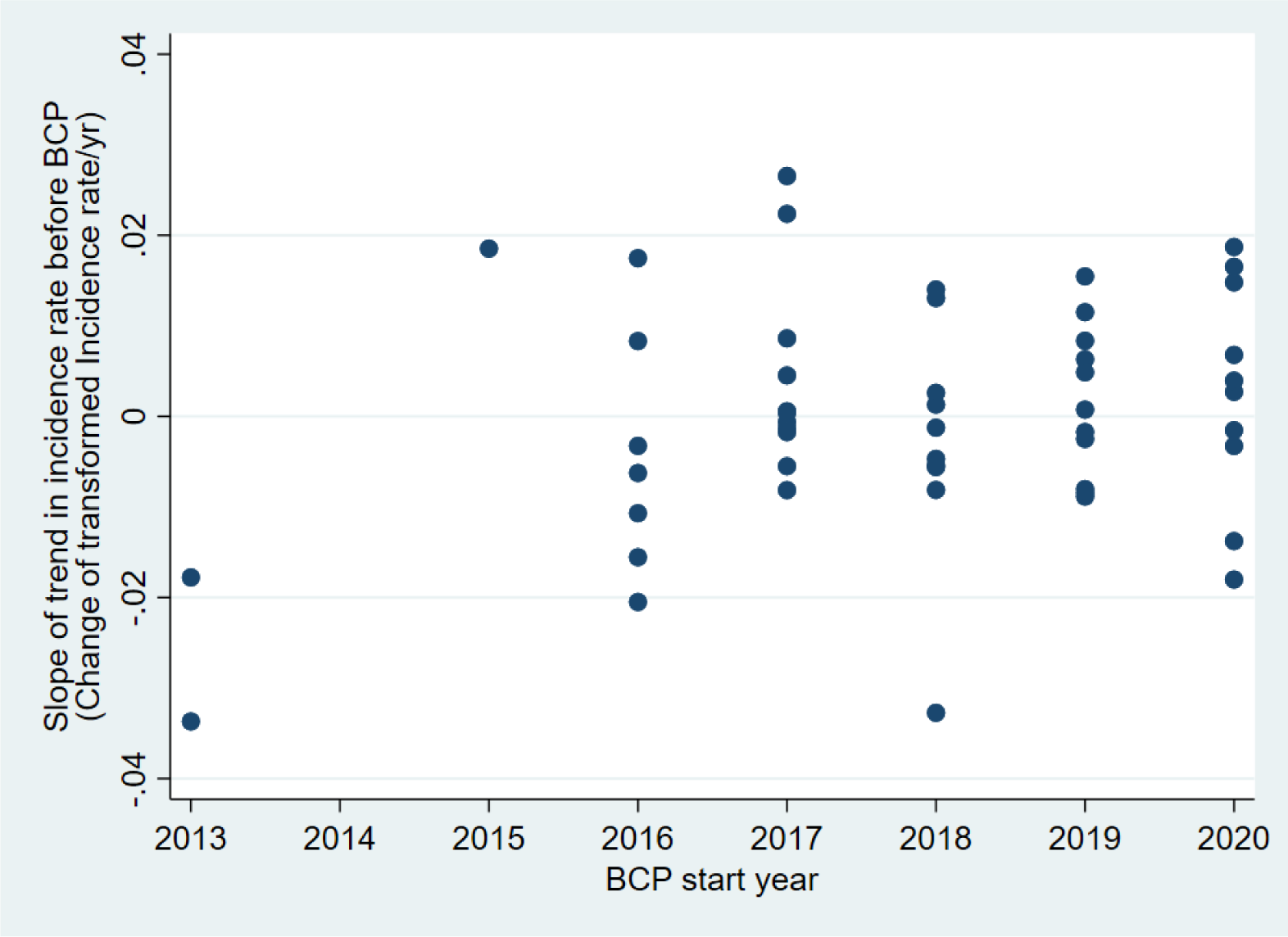
Relationship between trend in bovine TB herd incidence rate in individual areas before the local start of BCP and the year BCP started.

0.135 or 0.165. Thus, the strongest trends observed before BCP were up to the magnitude of the estimated effects of the BCP (56% reduction in the 4^th^ year). The slopes estimated for areas 1 and 2 had more negative values than most other areas. Their uncertainty may be relatively large because they were observed for just 4 years before the start of BCP, whereas all other areas were observed for at least 6 years.

### Comparison of primary and alternative analyses

The primary analysis was compared with analyses of alternative data and using a Poisson GLM to confirm the estimated effects of the BCP were not excessively dependent on the specific analysis applied. The estimated outcomes were similar when using intervention years and calendar years, with confidence intervals overlapping across most of their ranges (Table 3). The estimated reduction of the total incidence rate was substantial at 45%, although 10-12% less than the reduction of the OTFW incidence rate. Analysis using data on cohort herds instead of herds in existence (HIE) estimated smaller reductions associated with the BCP (Table 3). The TB incidence rate among cohort herds increases relative to HIE with time after or before the start of BCP, which suggests herds that persist in the cohort have a higher incidence rate than herds that are new or leave the cohort (Fig. A1 in Appendix).

**Table 3:** The BCP effect after the fourth year of controls for the alternative combinations of measures. Estimates are from a linear model of square root transformed incidence rate.

| Herds | Year | Incidents | Average Reduction (%) | 95% confidence interval |
| --- | --- | --- | --- | --- |
| HIE | Intervention | OTFW | 56 | 45 - 65 |
| HIE | Calendar | OTFW | 58 | 46 - 69 |
| HIE | Intervention | All | 45 | 35 - 54 |
| HIE | Calendar | All | 48 | 37 - 58 |
| Cohort | Intervention | OTFW | 48 | 36 - 59 |
| Cohort | Calendar | OTFW | 48 | 34 - 60 |
| Cohort | Intervention | All | 36 | 25 - 46 |
| Cohort | Calendar | All | 37 | 25 - 49 |

The estimates from using the Poisson regression method were close to the primary linear analysis presented above (Table 4). The largest difference between methods was when analysing the effect of BCP on total incidence rate in HIE, when the Poisson result was 5-6% lower than the primary analysis. A Jack-knife analysis to check the robustness of the statistical analysis found that partial estimates of the BCP effect from omitting individual areas were distributed around the estimate from all areas ^[27]^, with little evidence of bias (Fig. A2 in Appendix). In the Jack-knife analysis, responses of the primary analysis and the Poisson regression were correlated and had similar magnitude.

**Table 4:** The BCP effect after the fourth year of controls for the alternative combinations of measures. Estimates are from a Poisson regression model of counts.

| Herds | Year | Incidents | Average Reduction (%) | 95% confidence interval |
| --- | --- | --- | --- | --- |
| HIE | Intervention | OTFW | 55 | 47 - 62 |
| HIE | Calendar | OTFW | 57 | 48 - 65 |
| HIE | Intervention | All | 40 | 32 - 48 |
| HIE | Calendar | All | 42 | 32 - 51 |
| Cohort | Intervention | OTFW | 50 | 41 - 58 |
| Cohort | Calendar | OTFW | 51 | 40 - 60 |
| Cohort | Intervention | All | 33 | 24 - 42 |
| Cohort | Calendar | All | 34 | 22 - 44 |

## Discussion

The average effect associated with BCP controls estimated here was roughly consistent with previously reported effects of badger culling on TB in cattle from the BCP, RBCT and Ireland (Table 5). It was between the previous estimates from the BCP in Gloucestershire and Somerset. The effect reported from the RBCT was arguably lower. However, the RBCT began over 13 years before the BCP, when the TB epidemiological situation was different. Cattle movements may have made a greater contribution to transmission of bovine TB at the time of the RBCT. Restocking of cattle farms after the Foot and Mouth disease outbreak in 2001 was a contributory factor ^[28]^ and compulsory pre-movement TB testing of cattle did not start until 2006 ^[29]^. Higher numbers of infected cattle per breakdown and fewer controls on transfer of TB by livestock movements would have reduced the relative effect of transmission from badgers.

**Table 5:** The BCP effect reported from the primary analysis in this paper compared to previous published effects of widespread badger culling on TB incidence in cattle. RBCT is the randomized badger culling trial. Reductions are estimated relative to unculled equivalent sites. Gloucestershire, Somerset and Dorset are areas 1 - 3 in the BCP.

| Source | Timing | Estimated Reduction (%) | 95% confidence interval |
| --- | --- | --- | --- |
| BCP this study | Year after 4<sup>th</sup> cull | 56 | 43 - 67 |
| BCP Gloucestershire<sup>[21]</sup> | Year after 4<sup>th</sup> cull | 66 | 53 - 75 |
| BCP Somerset<sup>[21]</sup> | Year after 4<sup>th</sup> cull | 37 | 16 - 52 |
| BCP Dorset<sup>[21]</sup> | Year after 2<sup>nd</sup> cull | -10 | -58 - 23 |
| RBCT<sup>[16]</sup> | 4th cull to end of trial | 32 | 12 - 47 |
| RBCT<sup>[16]</sup> | 0-30 months after trial | 42 | 24 - 56 |
| Irish Four Counties<sup>[14]*</sup> | 1<sup>st</sup> cull to end of trial | 71 | 54 - 82 |

\* Effects are quoted from the cited publications, apart from the Irish 4 counties study, as explained in the text of the discussion.

The paper from the Irish Four Counties study in its Table 9 reported the change in TB incidence rate in cattle herds after badger culling as a coefficient of a log-hazard function ^[14]^. It presents coefficients for reference and removal areas in each of its four counties. As an analogous approach to the Difference in Differences analysis presented here, the difference between the removal area coefficient and the reference coefficient is an estimate of the coefficient for the effect of culling within each county. The reduction in Table 5 is from the exponential of the average estimated coefficient for the effect of culling across the four counties. The confidence interval is estimated from the standard error estimated from the variation among the four counties. The effect of badger culling reported from Ireland was higher than from the analysis reported here. However, the two confidence intervals overlap, and the badger populations and the epidemiology of bovine TB differ between England and Ireland ^[9,30]^.

Compared with previous estimates, this analysis is more precise on the timing of the effects associated with repeated badger culling (Fig. 4). The effect of badger culling is shown to increase each year for at least the first three years after the start of badger culling, with relatively little impact in the first year. Evaluations of the effect of control measures targeting wildlife reservoirs should take account of the timing of observations relative to the timing of interventions.

From April 2017, additional compulsory interferon-gamma testing of cattle was introduced to detect and remove infected cattle during TB incidents in areas that had been in the BCP for at least two years ^[20]^. So, the additional interferon-gamma testing was applied in all BCP areas, starting with the third intervention year in all but the first two areas. Otherwise, this testing was only compulsory for OTFW incidents in the Edge area. This additional testing aimed to supplement the effects of badger culling by reducing recurrence of infection in herds after the end of restrictions. The BCP effects in the first two intervention years took place before the additional testing started. Moreover, the additional testing only applied within herds already restricted after detection of TB by standard skin testing, which limited its influence in the third intervention year. However, this data analysis cannot explicitly distinguish the effects of the BCP’s component measures. The influence of additional gamma testing may be investigated by analysing the contribution of reduced recurrence of infection to the overall reduction of incidence rate. Further analysis comparing the effect of BCP controls in different areas may provide more specific evidence on the effects of badger culling.

Here the primary analysis was of OTFW incidence rate rather than the total incidence rate. As explained in the methods, OTFW incidence rate was expected to be the better surrogate measure of level of TB infection in cattle. The abrupt trends in OTFS incidence rate shown by Fig. 2, which are not matched by trends in OTFW incidence rate (Fig. 1), are more likely to be the result of changing surveillance than changes related to infection levels in cattle. Bovine TB surveillance in England is complex, with various changes during 2009-2021 ^[31]^. Moreover, reviews and metanalyses have suggested spatial temporal heterogeneity in the performance characteristics of bovine TB surveillance tests ^[32,33]^. Additionally, OTFS cases may be less indicative of an embedded local reservoir of infection with infected badger involvement ^[34]^. In practice, this study found a substantial reduction of total incidence rate including OTFS incidents, although less than of OTFW incidents only (Tables 3, 4). Similarly, the main conclusions of the analysis were not substantially changed by other variations of the data analysed nor by the method of analysis.

This analysis evaluates the outcome of a Government policy and not a controlled experimental trial. There are challenges to such analyses, including adequate adjustment for non-random variation and confounding factors. Choice of areas was by the farming industry subject to licence criteria, and developed over time. As a result, it was not possible to define all areas that would be included in the BCP at the start of the programme, nor to randomize the sequence in which the areas joined the BCP. The Difference in Differences analysis addressed this lack of randomization, adjusting for arbitrary differences in incidence rate between areas. BCP interventions in 49 areas started in five different years, with two more start years for the remaining three areas (Table 1). Thus, the analysis included many comparisons between years within areas, between pairs of areas while neither had started BCP, and between the same pairs of areas when they were at different stages in the BCP. There was evidence of trends in TB incidence rate specific to each area before the start of BCP, which were included in the model (Fig. 6, Table A1). Equivalent trends were obscured after the start of BCP by confounding with the effects of BCP and variation of those effects. A possible source of bias was for the potential trend in incidence rate without BCP interventions to differ between areas starting BCP at different times. However, the estimated trends before BCP started in each area were not associated with when BCP started (Fig. 7). Bias might also be detected by estimating the effect of BCP when it can be compared with an expected level. The analysis estimated near zero effect of BCP in its first calendar year, which matched prior expectations, consistent with little bias (Fig. 4b). In summary, the non-random starting sequence of BCP areas could have biased the estimated effect of BCP. However, there is no evidence of bias in the estimated effect, and we have not identified a feature of the starting sequence that would generate bias through the analysis applied.

A policy to control TB in cattle using badger culling must be bounded by ethical considerations ^[17]^. The current BCP is scheduled to finish within three years ^[35]^, although there may be some targeted intervention in the future in epidemiologically defined areas, where data demonstrate the level of risk from badger TB infection is of particular concern ^[12]^. Reducing risk of *M. bovis* transmission between cattle and badgers across large areas of England may increase the effectiveness of controls of other sources and pathways of infection for cattle. The current analysis and other work strongly suggest that reducing infection from the badger population reduces TB incidence rate in local cattle. Similar effects may be achieved or maintained by other measures, such as badger vaccination ^[36,37]^, fertility control ^[37]^ and biosecurity ^[38]^, although there is more evidence on the effect of culling than other options. Field studies in Ireland have shown that badger vaccination is a potential substitute for culling ^[39]^. However, to avoid unrealistic expectations, the delay reported here between starting controls and achieving results should be expected from any controls of bovine TB in wildlife, and may be longer for vaccination of wildlife ^[40]^.

In conclusion, in this study we report a substantial reduction in herd incidence rate of TB in cattle following introduction of the BCP, using a novel application of the econometrics method Difference in Differences for epidemiological analysis of a complex intervention policy over many years.

## Acknowledgements

We thank Phil Hogarth, Rachelle Avigad, Graham Smith, James McCormack, Pilar Romero, Ricardo De la Rua-Domenech, Nick Lyons and Christl Donnelly for comments on drafts of the manuscript. Funding for the project was provided by Defra (projects SE3131 and TBOM1500).

## APPENDIX

### Analysis of variance of factors affecting incidence rate

An analysis of variance was defined to closely match the constrained regression by selecting a suitable base area for the estimation of the slope parameter a_i_. Selecting area 41 to be the base area set a_41_ = 0, which generated estimates of a_i_ with average value 0.00083, which was a good approximation to the value zero defined by the constrained regression. Hence the sums of squares from the analysis of variance also described the constrained regression (Table A1). The model was dominated by the area effect; the model SS increased less than 50% with the inclusion of all other factors. Although the BCP effect was clearly significant and was the output of primary interest, it had little influence on the overall model fit. Thus, comparison of alternative data and methods of analysis and investigation of the model’s reliability had to focus on the estimates of BCP effect rather than diagnostics of overall model fit.

**Table A1:** Analysis of variance equivalent to the constrained regression used to estimate the effects of the Badger Control Policy. Sequential sums of squares (SS) show how the variance explained by the model increased as factors were added to the model in the order displayed. Partial SS show how the model was improved by adding each factor last. Mean squares (MS), F ratios and significance are calculated from the partial SS.

| Source | Sequential SS | Partial SS | df | MS | F | Prob>F |
| --- | --- | --- | --- | --- | --- | --- |
| Model | 4.439 |  | 118 | 0.0376 | 22.9 | 0 |
| Area effect | 2.992 | 1.623 | 51 | 0.0318 | 19.37 | 0 |
| Year effect | 0.484 | 0.0656 | 11 | 0.00596 | 3.63 | 0.0001 |
| Pre-BCP slope by area | 0.869 | 0.617 | 51 | 0.0121 | 7.37 | 0 |
| BCP effect | 0.094 | 0.0938 | 5 | 0.0188 | 11.42 | 0 |
| Residual | 0.830 |  | 505 | 0.00164 |  |  |
| Total | 5.269 |  | 623 | 0.00846 |  |  |

### TB incidence rate among herds in existence versus among cohort herds

**Figure A1:**
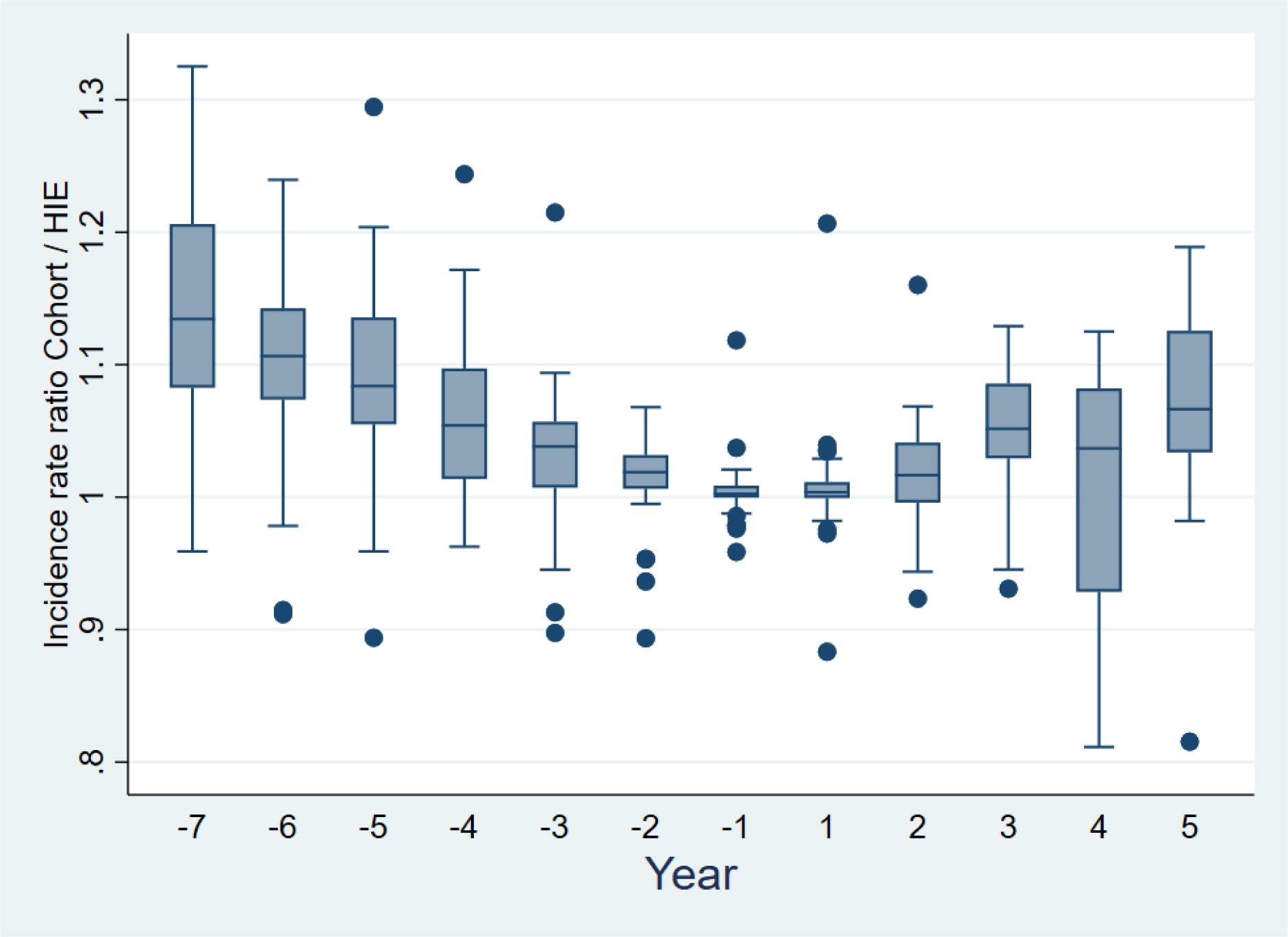
Box and whisker plot showing the median ratio of TB incidence rate among cohort herds divided by incidence rate among herds in existence (HIE) increased with time before or after the start of BCP interventions. In the box and whisker plot, the central horizontal line is the median. The box ranges from the lower quartile to the upper quartile. The whiskers show the full range of the data, unless outliers are present, which are shown individually as dots.

### Jack-knife resampling of estimates of BCP effects

A check of the robustness of a statistical model is to recalculate its critical outputs after removing individual observations. This check can reveal excessive dependence of model outputs on small portions of the data. Figure A2 shows estimates of the BCP effect in the fourth year recalculated after removing each of the 52 areas.

**Figure A2:**
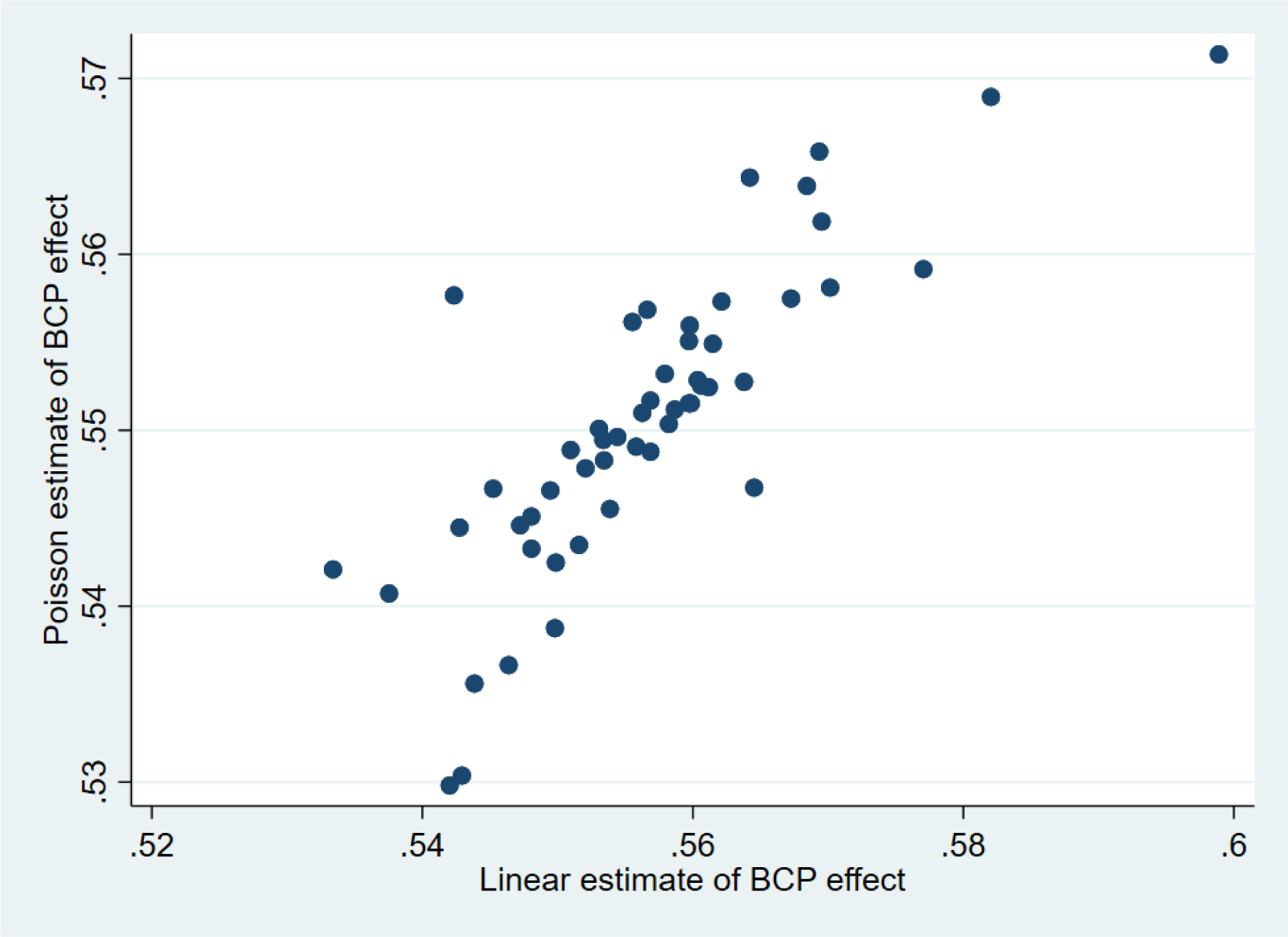
Plot of the partial estimates from a Jack-knife analysis in which each of the 52 areas was omitted in turn. The BCP effect in its fourth year estimated by the Poisson analysis is plotted against the estimates from the primary, linear analysis. The partial estimates are centred close to the estimates from the full analyses (0.5566 from the linear analysis and 0.5505 from the Poisson analysis).

## Notes

### Competing Interest Statement

The authors have declared no competing interest.

